# Combining artificial intelligence and local ecological knowledge to document the largest ever-counted Cape buffalo mega-herd

**DOI:** 10.1101/2025.11.26.689243

**Authors:** Emily Bennitt, Blair R. Costelloe, Benjamin Koger, Elodie Wielgus, Gaseitsiwe Masunga, Alexandre Caron

**Author notes:** **Correspondence:**; Okavango Research Institute, Shorobe Road, Sexaxa, Maun, Botswana. Joint first authors (Emily Bennitt, Blair R. Costelloe).

## Abstract

Large aggregations of wild mammals are declining globally, sometimes before they can be scientifically documented and their ecological value understood, although non-scientists may be aware of these phenomena. Video recording of wildlife by non-academics is becoming more frequent with increasing activities of humans in natural habitats. This opportunistically-collected material can document rare or ecologically-important events, such as large aggregations, and thus provide potentially valuable data for ecologists and conservationists. We used footage of a large herd of Cape buffalo recorded by a wildlife film producer in the Mababe Depression, northern Botswana, and applied automated detection and tracking techniques to count individuals as they moved across the video frame. To complement this demographic snapshot, we accessed local ecological knowledge from stakeholders to provide contextual information for a more complete understanding of the ecosystem processes potentially driving population movement and trends. Manual review of the automated count resulted in a minimum group size of 3676 buffalo and a total estimated herd size of 4131 (range 3828-4446), which matched herd size estimates provided by local stakeholders. To our knowledge, this is the largest documented group of Cape buffalo, which we identify as a mega-herd and recommend research into the costs, benefits and ecological consequences of forming such large groups. This study represents an opportunistic collaboration among wildlife filmmakers, computer scientists, lay experts and ecologists, and highlights the value of combining contributions from different fields to generate information that can be used in conservation practice.

## INTRODUCTION

Historically, large African herbivores came together in vast aggregations, congregating on resources or undertaking treks across landscapes (OwenSmith et al., 2020). The formation of large herbivore groups can confer costs, such as exposure to disease (Wielgus et al., 2021), and benefits, such as safety from predation (Bond et al., 2019). Large groupings still occur in some ecosystems, such as blue wildebeest (*Connochaetes taurinus*) in the Serengeti, Tanzania (Mduma et al., 1999), or white-eared kob (*Kobus kob leucotis*) in South Sudan and Ethiopia (Schapira et al., 2017). However, unfragmented landscapes that allow movement and congregation of herbivores are becoming rarer with anthropogenic developments and increasing demand for dwindling resources and many herbivores populations globally are declining (Kauffman et al., 2021).

Unfortunately, few of these movement and aggregation occurrences are well characterized and some are probably yet unknown (Kauffman et al., 2021). The detailed documentation of wildlife phenomena such as large aggregations of animals requires information on the number of those animals as well as their ecological role and the factors leading to group formation and dissolution (Kiffner et al., 2020). Often such information can only be garnered by scientists through years of intensive and detailed study, using GPS-enabled collars (OwenSmith et al., 2020) or camera traps (Cordier et al., 2022). However, some wildlife professionals, while not scientists, routinely observe and, in some cases, document wildlife aggregations and thereby gain insight into the drivers of these phenomena. Those professionals, including wildlife film-makers, professional guides and local communities in wildlife areas, may have observed the functioning of the ecosystem around such aggregations over numerous seasons, leading to a fount of anecdotal knowledge that should not be ignored based on a lack of academic rigour (Adade Williams et al., 2020; Bennett, 2016). Using material and knowledge from nonacademic actors can provide opportunities for scientists to fill knowledge gaps left by a lack of funding, time and capacity to collect scientific data (Tuia et al., 2022). However, the integration of local ecological knowledge (LEK) into scientific publications requires interdisciplinarity and, sometimes, unconventional approaches and methods (Dickinson et al., 2010).

Videos captured by wildlife film-makers can document unusual behaviours or sightings. Often, impressive aggregations are targeted by film-makers who use drones to capture footage which, while not designed for accurate population counts, can be used to secure estimates of group size (Hodgson et al., 2018; Pollock et al., 2023). Automated deep-learning based processing has proven successful at efficiently counting animals in aerial images, but accurately extrapolating these counts to population estimates can be complicated by the risk of double counting animals when survey images overlap in space (Delplanque et al., 2022; Eikelboom et al., 2019; Kellenberger et al., 2018; Torney et al., 2019). An extreme example of such a case is video-based observation of mobile animals filmed at close range. Here, automated processing tasks become more complex since animals must not only be detected but also tracked across many video frames to avoid over-counting (Koger, Hurme, et al., 2023).

Population counts provide a demographic snapshot of the number of individuals but offer minimal insight into the ecological processes or ecosystem functions that support or are facilitated by animal populations (Kiffner et al., 2020): counts convey limited information about movement patterns, possible threats to the population, including at animal-human interfaces, drivers of distribution or how that population functions within the ecosystem. Given the wealth of media, including photographs and footage, being recorded by non-scientists, it is important to develop tools to harness valuable data being collected ad-hoc across the world, and integrate knowledge from non-scientist professionals and indigenous communities. Multi-pronged approaches that combine count data with other information are vital to understanding the link between environmental conditions and population dynamics (Marsh & Trenham, 2008), thereby increasing the reliability of population viability predictions in areas predicted to be hit hard by climate change, such as northern Botswana (Engelbrecht et al., 2015).

The northern Botswana conservation zone represents a large, ecologically functional and connected landscape that hosts the longest plains zebra (*Equus quagga*) migration in the world (Naidoo et al., 2016) and allows adaptive movements between seasonally productive landscapes. Herbivores in northern Botswana tend to move between areas with permanent water during the dry season and more wooded and sandy habitats during the rainy season, when these habitats produce nutrient-rich annual grasses (Bennitt, Hubel, et al., 2019). The Mababe Depression (MD), part of an ancient lake system in northern Botswana (Burrough et al., 2009), is fed during the dry season by water flowing from the Khwai River at the southern edge of the MD. The volume of water following this route appears to have increased in recent years (Bennitt, 2024, pers.obs.), likely in response to a combination of tectonic shifts and human activities. During the dry season, Cape buffalo (*Syncerus caffer caffer*) have been observed congregating in large numbers around the marshy habitat at the south of the MD. Such large seasonal congregations of buffalo, which represent several mixed herds coming together, have been defined as mega-herds, and most mixed buffalo herds number less than 1,000 individuals (Caron et al., 2023). Stakeholders operating in the area have been reporting observations of these aggregations for over a decade. Aerial surveys conducted in 2015 and 2018 estimated numbers of buffalo in the MD at 2788 (Chase et al., 2015)) and 3498 (Chase et al., 2018), respectively. Accurate counts of mobile animals occurring in such large numbers can be very difficult, even with a series of still images or aerial footage (Koger, Hurme, et al., 2023). However, recent advances in artificial intelligence have supported the development of algorithms to detect, track and count individual animals moving through frames of video footage (Koger, Hurme, et al., 2023; Rathore et al., 2023).

In 2022, a team from Silverback Productions (Bristol, UK) for Netflix (Los Gatos, California, USA) captured drone footage of a very large buffalo group. They contacted academics specialized in African buffalo ecology (from the authorship) to help them estimate the herd size. Recognizing the exceptional size of this herd based on visual examination of the video, we decided to make the most of this opportunistic media material using a mixed method. We developed a deep learning-assisted procedure for counting every buffalo captured in the video, allowing the most accurate count to date of this mega-herd. To better understand the dynamics leading to the formation of the mega-herd and its impacts on the ecosystem of the MD, we gathered LEK through interviews with knowledgeable stakeholders living and/or operating in the area. Specifically, we wanted information that would allow us to (i) determine the size of the Mababe buffalo herd, (ii) define the time period when buffalo congregate in the MD, and (iii) investigate the socio-ecological role of buffalo in the MD.

## METHODS

### Study site

The MD straddles the border between the Chobe National Park and a wildlife management area, NG41, that supports trophy hunting and photographic tourism (Fig. 1). As a consequence of being part of the Makgadikgadi Paleolake that extended over 66,000 km^2^ at its largest extent (Burrough et al., 2009), the MD contains nutrient-rich lacustrine soils that support productive grasses attracting multiple species of herbivore (Fynn et al., 2014). In particular, soils and vegetation in the southern part of the MD contain high phosphorus, potassium and calcium levels (Sianga & Fynn, 2017). The Mababe community has a settlement at the southern end of the MD, where they engage in activities and livelihoods based on natural resources, such as tourism, veldt product harvesting and fishing.

**Figure 1.**
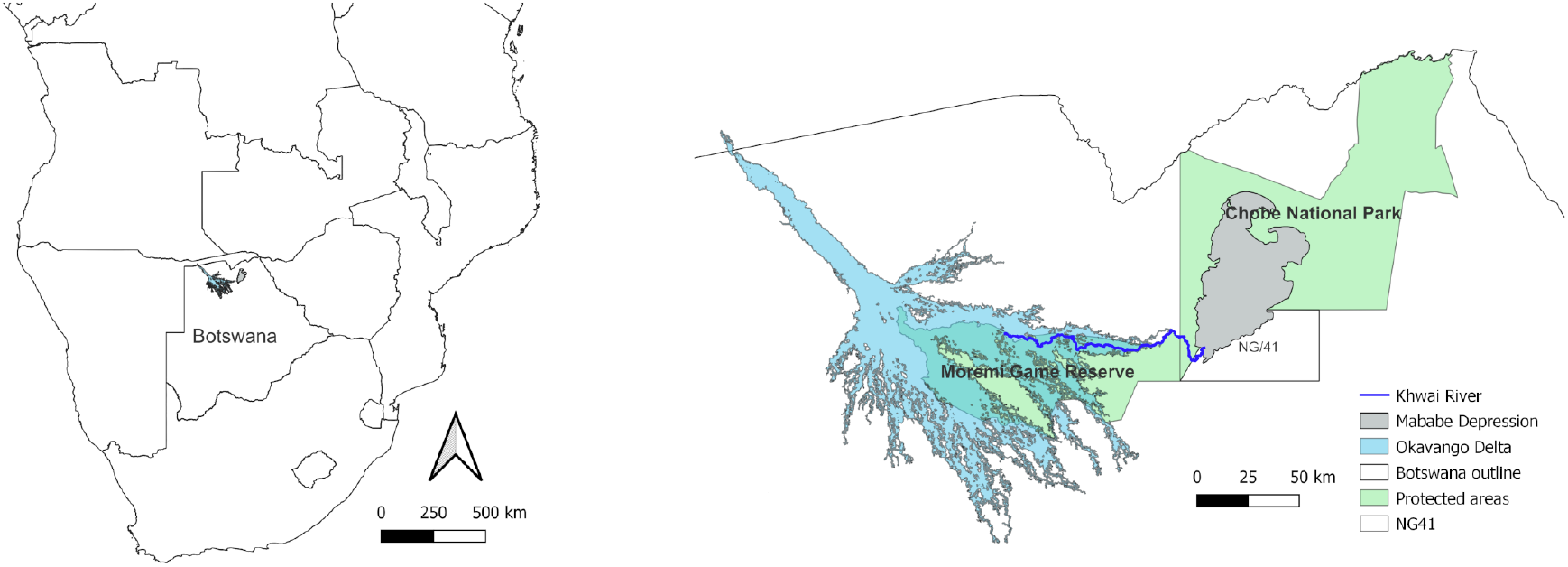
Map of study area, highlighting the Mababe Depression in northern Botswana, where several thousand Cape buffalo congregate during the dry season.

### Buffalo count

We counted buffalo in a video of the mega-herd that was provided by Silverback Productions in collaboration with Netflix. The buffalo were filmed from 09:39 for 23 minutes on the 26th September 2022 using a DJI Mavic 3 Pro Cine (Nanshan, Shenzhen, China) drone fitted with a 4/3 CMOS Hasselblad (Gothenburg, Sweden) camera. The drone hovered in a stationary position approximately 100 m above the herd and filmed at an oblique angle as the herd, moving cohesively in one direction, passed under the camera (Fig. 2). The film-maker aimed to capture natural behaviour so minimized the impact of their presence (and that of their equipment) by launching the drone from over 200 m from the edge of the buffalo herd. In general, African herbivores are only disturbed by a drone when it is <60 m above ground level and <100 m horizontal distance from them (Bennitt, BartlamBrooks, et al., 2019). The footage shows no indication of disturbance to the buffalo caused by the presence of the drone, such as running or looking up.

**Figure 2.**
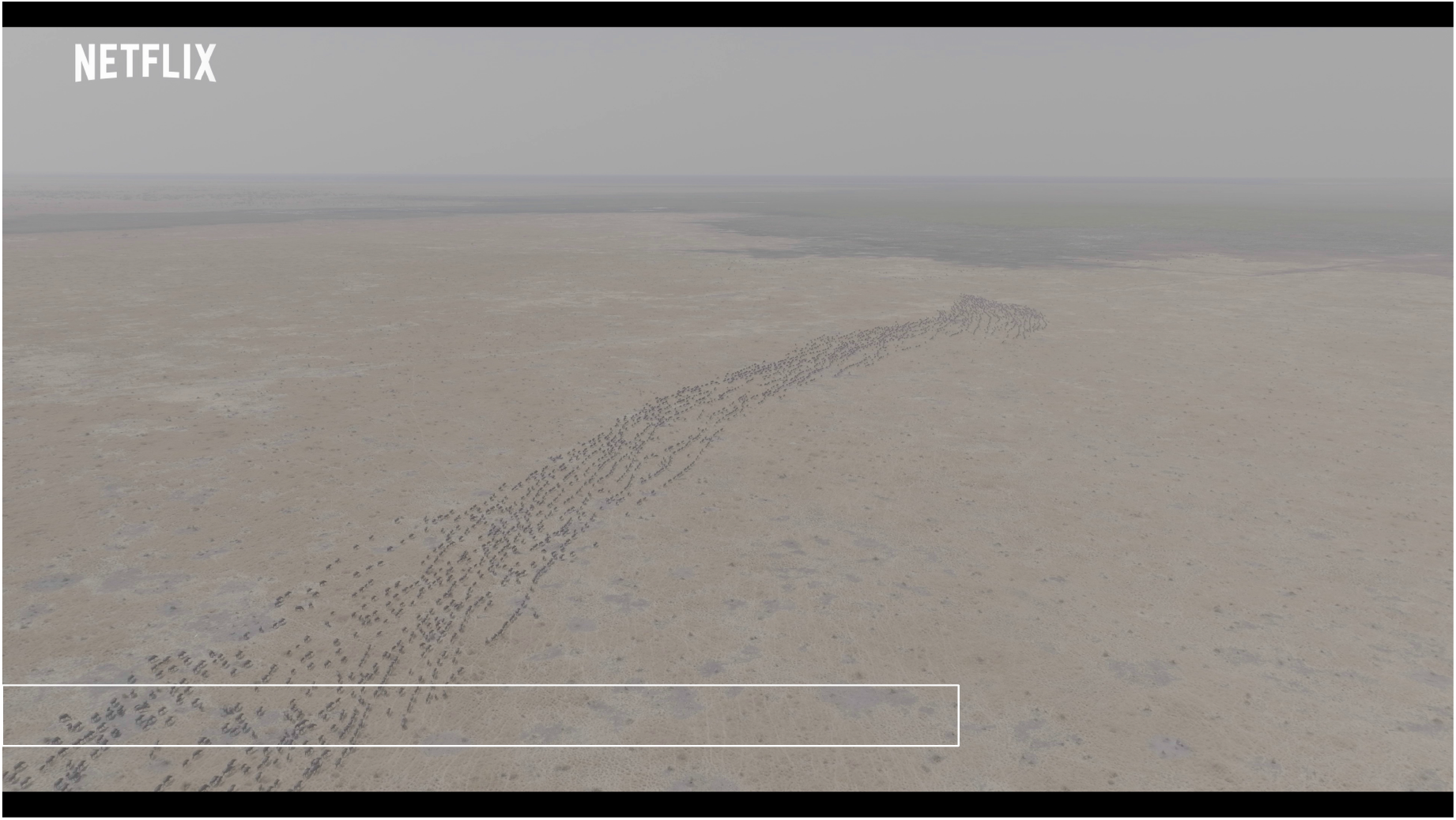
A still frame from the video provided by Netflix. The buffalo herd is moving away from the camera, toward the horizon. The white rectangle shows the area of our analysis, where we performed automated detection and tracking and manual count correction.

We used a deep learning-assisted approach, detailed below, to count the animals in the video: we first used methods adapted from Koger, Deshpande, et al., 2023 to detect and track individual buffalo in the video. We then systematically reviewed the tracked video using an approach modified from Koger, Hurme, et al., 2023 to manually count buffalo that the detection and tracking procedure missed. We summed the number of automatically-detected tracks and the manually counted animals to calculate the total number of animals in the video, adjusting for the detection probability and precision of the human reviewers. The video ends while animals are still entering the video frame. Therefore, we fit a regression model to forecast the rate of buffalo passage after the video ends and used this model to estimate the number of animals missed due to early termination of the video. We provide the data and Python code necessary to reproduce our results in an open-access data repository and on GitHub.

### Buffalo detection and tracking

The video clip was provided to us in 4K resolution (3840 x 2160 pixels). For object detection, we cropped out a 2530 x 160 pixels area near the bottom of the video frame, where the animals were closest to the camera and thus appeared the largest (Fig. 2). All buffalo passed through this area moving away from the camera, from the bottom of the frame to the top. No buffalo moved in the opposite direction. To ease the processing load, we only processed every third frame, effectively down-sampling the video from 24 to 8 frames per second (fps). We then used these processed frames to create videos for human review at a framerate of 30 fps: thus, the total video length was reduced from 22 minutes and 58 seconds to 6 minutes and 8 seconds. This effectively resulted in a video that was sped up 3.75 times, which eased the manual review process (see below) by reducing the total amount of footage that human reviewers needed to watch.

To detect buffalo in the cropped and down-sampled video, we used the Detectron 2 API (Wu et al., 2019) and code modified from Koger, Deshpande, et al., 2023 to train a FasterRCNN (Ren et al., 2017) object detection model with a resnet-50-fpn backbone (Lin et al., 2018). To build a set of annotations for model training, we randomly extracted still frames from the cropped video and extracted a randomly-located 100 x 100 pixel crop from each extracted frame. We used Labelbox (https://labelbox.com) to draw bounding box annotations around each buffalo in these cropped images. After initial training efforts resulted in detection failures where buffalo were in dense clusters, we extracted additional frames that included examples of this scenario, and from these extracted randomly-located 160×160 pixel crops, which we again annotated in Labelbox. Annotated cropped images were then randomly assigned to either the training or validation sets (Table 1). We modified the default Detectron 2 parameters to include images with no annotations in the training set; this helped eliminate instances where the model mistakenly identified tufts of grass as buffalo. While we evaluated the models raw detection performance on the images in the validation set (see Supplement for details), the process of evaluating the accuracy of the actual herd estimate these detections helped create followed a separate process described below.

**Table 1.** Number of cropped frames and annotations used to train and validate the detection model used to count a buffalo mega-herd from drone footage taken in the Mababe Depression, northern Botswana, in 2022.

|  | Cropped frames | Total buffalo annotations |
| --- | --- | --- |
| **Training set** | 366<br>(318 100x100 crops<br>+ 48 160x160 crops) | 1876 |
| **Validation set** | 92<br>(80 100x100 crops<br>+ 12 160x160 crops) | 484 |

After processing the video with the trained model, we connected detections across frames to generate movement trajectories, or tracks, for individual buffalo, again using code modified from Koger, Deshpande, et al., 2023 (Supplementary Video 1, doi xxxx). We discarded very low-quality detections, using only detections with confidence scores greater than 0.2 for track generation. This process generated 4868 movement trajectories. However, this number is not an accurate indication of the number of animals in the video since imperfections in buffalo detection resulted in imperfections in the trajectories. Some trajectories ended prematurely (before the buffalo left the frame) and then later restarted, resulting in multiple tracks for a single buffalo. In other instances, missed detections resulted in tracks that jumped between multiple individuals. These errors caused some buffalo to be counted multiple times and others to be missed entirely. Therefore, human review was necessary to correct the count generated by the automated tracking procedure.

### Manual review process

To correct the count generated by the tracking procedure, we used a manual review approach similar to that used by Koger, Hurme, et al., 2023 to count flying bats emerging from a roost. We designed a task in which human reviewers counted the number of buffalo that crossed a horizontal count line without being detected by the automated tracking algorithm.

We first drew a count line horizontally across the middle of the cropped buffalo video and added a buffer of 5 pixels on either side. We then created a version of the cropped video in which every track was represented by a colored dot corresponding to the centroid of the bounding boxes predicted by the detection model. This dot flashed with a yellow circle when it entered the buffer zone and crossed the count line (Supplementary Video 2, doi xxx). From the original 4868 movement trajectories, we retained only trajectories that started below and ended above the buffer, thereby eliminating partial tracks that either did not cross the count line or started or ended very close to the count line. This effectively eliminated instances of double-counting (see Results) and simplified the manual counting task. The total number of retained tracks was 2972.

It was very difficult for human reviewers to monitor the entire count line at once. Therefore, to make the task more manageable, we divided the video spatially into 5 horizontal sections (Fig. 3) and made a cropped video of each section. In these cropped videos, we added two red vertical lines to divide the count line into thirds. Reviewers could then focus on each third of the video separately, which further simplified the counting task. We divided these cropped videos temporally into 20-second segments (Supplementary Video 3, doi xxxx), resulting in a total of 95 video clips. Reviewers watched the clips and recorded, for each third of the video area (left, center and right), 1) the number of buffalo that were not counted by the tracking procedure (i.e., crossed the line without a colored dot), 2) the number of buffalo that were double-counted (i.e., had more than one colored dot associated with them) and 3) the number of non-buffalo objects that were marked with a colored dot. Each clip was reviewed independently by 3 reviewers (BRC, AC and EW). If all three reviewers agreed on the count for a given clip, that count was accepted as accurate. Clips for which reviewers disagreed (n= 34) underwent a second round of review. During this second round of review, a single reviewer (BRC) repeated the counting procedure for each of the 34 disputed clips and individually identified each missed buffalo, for example by noting the time stamp at which it crossed the count line, characteristics of the animal (e.g. calf or adult), and the color of neighbors tracking dots. The reviewer also noted instances where buffalo crossed the line very close to the edge of the cropped frame or near the start or end of a clip to make sure that individuals were not missed or double-counted across spatially or temporally adjacent clips.

**Figure 3.**
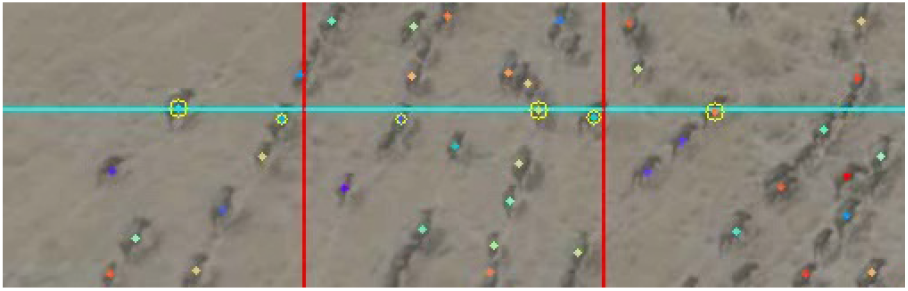
Astill frame from a video clip used in the manual count correction task. The vertical red lines ease the counting task by allowing the reviewer to concentrate on one third of the video frame at a time. Trajectories created by the automated detection and tracking process are depicted as colored dots overlaid on individual animals. As the trajectories enter the buffer zone around the cyan count line, they gain a yellow outline that grows in size as the trajectory crosses the count line, generating a yellow flash. Reviewers were asked to determine how many animals were missed by the detection and tracking process by counting the number of animals that crossed the line without a colored dot. For the full example video, see Supplementary Video 3.

We used a double-observer method (Nichols et al., 2000) to assess the accuracy of the second round of review. Eight randomly-selected clips were independently reviewed by an additional reviewer (BK), who also individually identified each missed buffalo. For each clip, the reviewers were randomly assigned as either the primary or secondary observer. We then compared the counts for each clip and identified how many individual buffalo were missed by the primary reviewer but found by the secondary reviewer. We then estimated BRCs detection probability using the following formula (Nichols et al., 2000):

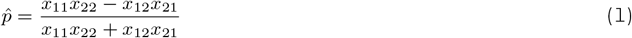

where *x*_*ij*_ is the number of individuals counted by observer *i* (*i* = BRC, BK) in clips where observer *j* (*j* = BRC, BK) was the primary observer. Here, *x* for the primary observer is the number of buffalo detected by the primary observer, whereas *x* for the secondary observer is the number of buffalo that were detected by the secondary observer but missed by the primary observer. This method allowed us to quantify BRCs detection probability, but did not allow us to quantify the probability of double-counting (false positives). To estimate the impact of double-counting, we calculated precision using the following formula:

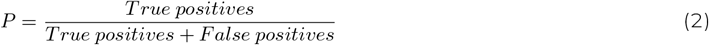

True positives and false positives were defined by reviewing and comparing the counts of BRC and BK in the subset of 8 clips that they both reviewed.

We calculated our herd count as:

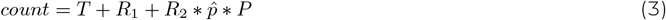

where *T* is the number of retained automatically-generated trajectories, *R*_1_ is the number of missed animals counted in the clips with no discrepancies in the first round of manual review, *R*_2_ is the number of missed animals counted by BRC in the second round of manual review, and 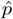 and *P* are BRCs detection probability and precision, respectively.

### Estimating the number of buffalo missed due to video ending prematurely

The video ends while buffalo are still entering the frame, suggesting that the herd size is larger than the number of individuals we are able to directly count, although the rate of buffalo moving through the frame and the width and density of the buffalo column suggests that the herd is tapering off. To estimate the number of animals that would have passed the count line if the video continued, we first plotted the count data as a time series. Using a range of bin sizes (2, 5, and 10 seconds), we calculated the number of automatically-generated trajectories that crossed the count line during each bin. We then summed the number of manually-detected buffalo for each 20 second video segment and divided this number by 10, 4 or 2 (depending on the bin size) and added this number to each bin corresponding to the range of the video segment. We then rounded the total count number for each bin to the nearest integer for modeling.

We fit a Bayesian Poisson regression model with a Gaussian-shaped time-varying intensity function to each time series. These models assume that the distribution of buffalo passage rates over time is bell shaped, with a slow start, a single peak and a gradual decline. However, they allow for uncertainty in the timing and intensity of the peak. We used this model to forecast the passage rates of buffalo in the 500 seconds after the video ends. Note that here 500 seconds refers to the time scale of the video, which is 3.75 times faster than real time due to our resampling of the frame rate (see Buffalo detection and tracking). Thus, we forecast buffalo crossing rates 1875 seconds ( 31 minutes) into the future in real time.

Mathematically, our models curve asymptotically approaches zero over time, but will never intersect zero; that is, the model will never predict an end to the herd (i.e. the condition where no buffalo are crossing the line). A protracted decline in buffalo passage rate is biologically unrealistic, as a long series of increasingly distant stragglers would be highly vulnerable to predation in this system. Therefore, we assumed for our analysis that the herd would end after the passage rate dropped below a given threshold. We defined thresholds of 1 buffalo crossing every 5, 10, and 20 seconds in real time (or 0.2, 0.1 and 0.05 buffalo per second, respectively). In the original video, it takes a single buffalo approximately 2.5 seconds to fully cross the count line. Using this crossing speed, we can estimate the spacing of individuals in body-lengths for each threshold duration using the following formula:

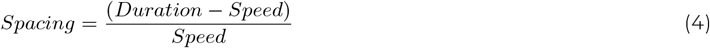

Thus, a passage rate of 1 buffalo every 5 seconds represents a single column of buffalo spaced approximately 1 body length apart while 1 buffalo crossing every 20 seconds represents a single column of buffalo spaced 7 body lengths apart. To estimate the number of buffalo missed, we assumed that the herd ends when the upper 95% confidence interval of our predicted passage rate drops below the threshold. We then calculated the cumulative number of buffalo expected to pass during the time between the end of the video and this projected herd end point.

### Local ecological knowledge

Contextual LEK on the Mababe mega-herd was sought from stakeholders living and working in the area, including safari operators, guides, filmmakers and local community members. All informants consented to be interviewed and, where relevant, signed consent forms allowing them to be cited in this manuscript. No personal information was gathered and the questions asked related only to the Mababe Cape buffalo mega-herd. The concession manager and the filmmaker responsible for the footage were the first informants, and they directed us towards other knowledgeable stakeholders. We initiated a community meeting to seek information from the Mababe community. We asked each informant about their knowledge of the existence of a large buffalo herd in the area, the estimated size of the mega-herd and whether it has changed over time, when the buffalo congregate, their movement patterns (including where they go outside of the dry season), and the role that they play within the MD ecosystem.

## RESULTS

### Buffalo count

Our detection model had an average precision of 0.868 (see Supplement for performance details). The automated tracking procedure generated 4868 trajectories. After eliminating partial tracks and those that did not fully traverse the buffer zone around the count line, we retained 2972 tracks. In the first round of manual review, the three human reviewers independently identified between 672 and 688 missed buffalo, zero instances of double-counting, and 1 non-buffalo object (a zebra). All three reviewers agreed on counts for 61 video clips, in which a total of 30 missed animals were counted. The remaining 34 clips underwent the second round of review, in which BRC counted 681 missed animals. Eight of these clips were also reviewed by BK in order to assess the accuracy of BRCs count. In these 8 clips, BRC counted a total of 103 missed animals and BK counted 99. In the 4 clips where BRC was the primary observer, she counted 41 missed buffalo and BK found 0 additional buffalo; in the 4 clips where BK was the primary observer, he counted 58 missed buffalo and BRC found an additional 3 missed buffalo (Table 2). Based on this analysis, BRC had an estimated detection probability of 1.0:

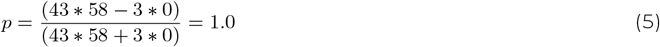

**Table 2.** Results of the double-observer procedure. Each reviewer served as the primary observer for 4 randomly-selected clips. The primary count is the number of buffalo that were missed by the tracking algorithm but detected by the primary observer. The secondary count is the number of buffalo that were missed by the tracking algorithm and the primary observer, but detected by the secondary observer.

|  | **Primary observer** |  |
| --- | --- | --- |
|  | **BRC**<br>(*n*=4) | **BK**<br>(*n*=4) |
| **Primary count** | 41 | 58 |
| **Secondary count** | 0 | 3 |

However, one buffalo counted by BRC was actually a double count. We therefore calculated the precision of BRCs count in addition to the detection probability:

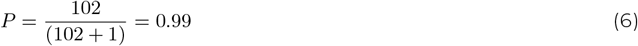

We calculate the total number of buffalo present in the video as:

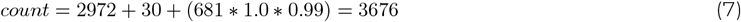

### Forecasting missed buffalo

Using a bin size of 5 seconds and assuming that the herd ends when the upper 95% confidence interval drops below 1 buffalo crossing every 10 seconds, we estimate that the herd continued for an additional 23.75 minutes (in real time) after the video ended (Figure 4). In this time, we estimate an additional 455 (range: 152-770) buffalo would have crossed the line. Thus, we estimate the total herd size as 4131 (range 3828-4446). These estimates were similar across the range of bin sizes and thresholds we considered (see Supplementary Materials for full details).

**Figure 4.**
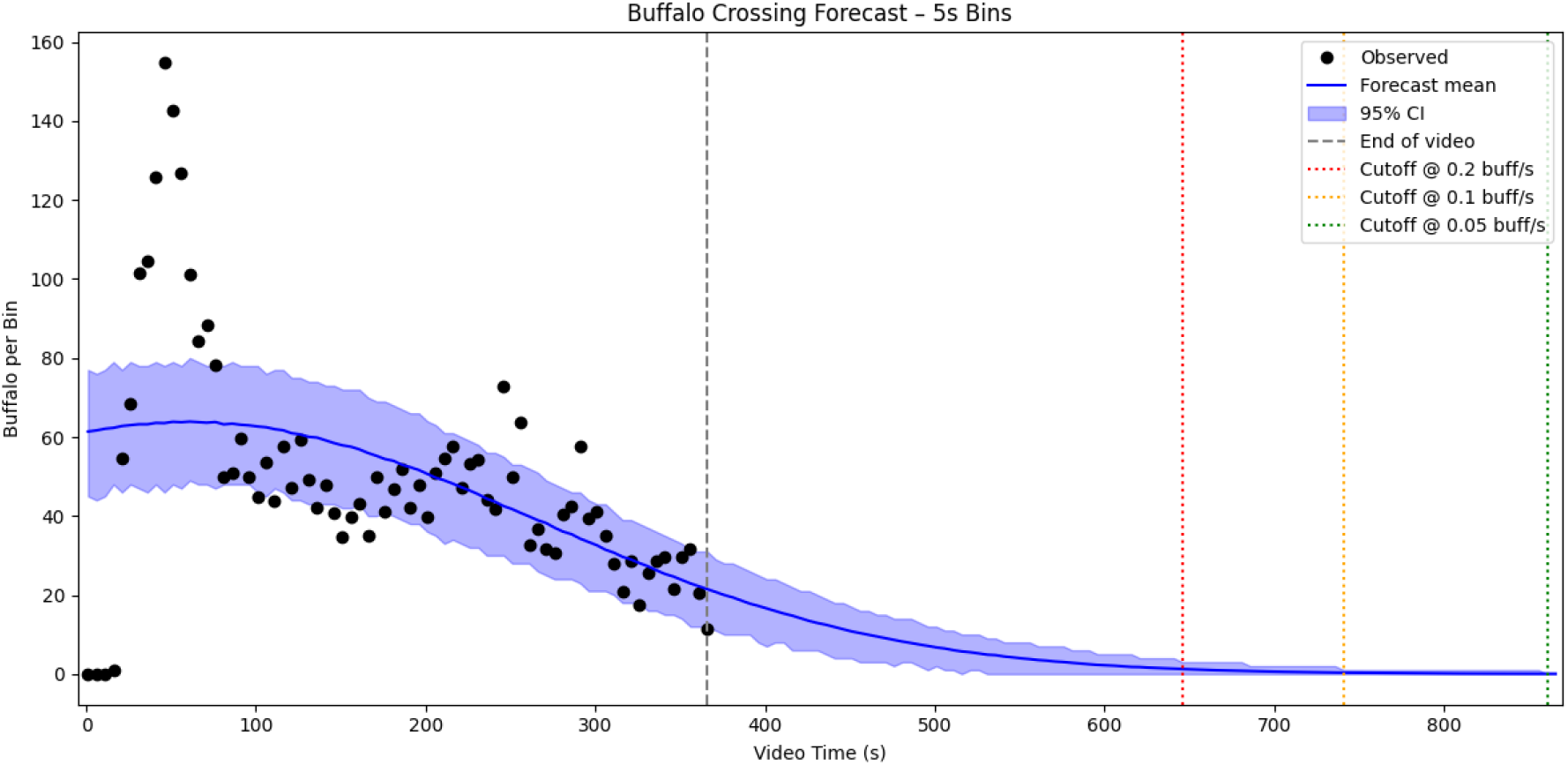
Atime series of the buffalo count data with fitted regression model. Data point represent combined automated and manual counts, binned into 5 second video segments. The 95% confidence intervals are generated by drawing samples from the posterior distribution of the models parameters. Vertical lines represent the end point of the video (grey), and the cut-offs used to estimate the end point of the herd (red, yellow, green), which are determined as the point at which the upper confidence interval drops below a given rate of buffalo crossings. Estimates based on a 5 second bin and a cutoff rate of 0.1 buffalo crossing per second are reported in the main text. Plots and results for additional time bins are provided in the Supplementary Materials.

### Local ecological knowledge

We collected information from one concession manager, two filmmakers and one professional guide, who was a member of the Mababe community, and we held a community meeting (kgotla) in Mababe village, which was attended by approximately 25 community members. Stakeholders from the wildlife industry apparently had more information about the mega-herd than community members, even though the latter were living full-time in Mababe, whereas the former were more transient in the area.

Stakeholder estimates for the mega-herd ranged from 3,000 to 4,000, which correlated closely with our video-based count. One stakeholder stated that other groups had also been observed in the area, for a total buffalo population up to 7-8,000 in the MD (Calitz, 2023, pers. com.). Most stakeholders from the wildlife industry reported a fluid population, with herds joining and leaving the mega-herd over the course of the season. Surprisingly, the community members had very little knowledge of the mega-herd and said that they were not able to count buffalo as these were usually in thick vegetation. Most comments from community members referred to buffalo being seen in and around the village, necessitating precautions to avoid potentially negative interactions. The lead escort guide for the community first heard of the mega-herd in 2019 but has never physically seen it and has not heard of it since (Kebuelemang, 2023, pers. com.). No stakeholders perceived a decrease in the population, and some indicated that there may be more buffalo coming into the MD with increasing water from the Khwai River since 2019 (Bestelink, 2023, pers. com.).

All stakeholders with knowledge of the mega-herd agreed that it was present in the MD over the dry season, from August to October, and absent during the rainy season, possibly because the soil in the MD becomes very waterlogged and difficult to walk on for large animals during the rainy season (Bestelink, 2023, pers. com.). During the rainy season, buffalo leave the MD and views differed as to whether they remain in the surrounding woodland area or move further afield, to the north or west. Productive grasses in the MD were viewed as a main attractant for buffalo, which were suggested to impose a heavy grazing pressure with the possibility of facilitation for smaller herbivores (Calitz, 2023, pers. com.). Large aggregations of buffalo in relatively open habitat were thought to be attracting high densities of male lions, with 18 different individuals recorded in 2022 and 10 buffalo kills documented in 21 days (Calitz, 2023, pers. com.). Predation pressure from lion hunts, which was mainly focused on weaker individuals (Bestelink, 2023, pers. com.), was reported to cause buffalo movement and drive splitting of the mega-herd.

## DISCUSSION

Accurate counts obtained through machine learning provide demographic information that needs to be paired with information on wildlife behaviour, movement patterns and ecology to create a comprehensive understanding of the factors that may affect population dynamics (Johnson et al., 2010), bringing together the expertise of data scientists and the experience and knowledge of field ecological experts (Tuia et al., 2022). In this study, skills and expertise from multiple arenas were harnessed to reach a common goal. Footage collected by wildlife filmmakers provided the required material for specialists in computer vision to develop and adapt algorithms to count a buffalo mega-herd. Researchers with a background in buffalo ecology helped to validate the count and we complemented the approach with LEK to understand factors driving the formation of the mega-herd and its broader impacts. Harnessing technological advances in video processing with tracking algorithms allowed us to produce an accurate count of a buffalo mega-herd that can be repeated over time to monitor population changes. Although information obtained by non-scientific methods can be inaccurate (Moqanaki et al., 2018), this count corresponded closely to estimates provided by local stakeholders, indicating that their understanding of ecology of the Mababe mega-herd is also likely to be reliable and highlighting the importance of incorporating LEK into conservation (Fazey et al., 2006).

We employed an interdisciplinary approach to maximise the use of data collected opportunistically and non-academically to report a previously undocumented natural phenomenon, the Mababe Cape buffalo megaherd, which includes a minimum of 3676 animals, making it the largest buffalo herd on record. In our estimate we used a conservative predictive model, which assumes only a single peak in buffalo crossings. However, we cannot rule out the possibility that additional peaks occurred after the end of the video, in which case the total herd size could be much larger than we estimate. The numbers we report should thus be taken as a conservative estimate of herd size.

The opportunistic co-production of science by a wide array of stakeholders provides valuable insights contributing to scientific knowledge in the global context of habitat fragmentation and gradual disappearance of large-scale ecological phenomena. Beyond gaining a better understanding of this important buffalo population, we demonstrated the ability of machine learning-assisted tools to accurately process opportunistic video footage of large transient animal aggregations. Deep learning methods may be time-consuming to develop and train, but they can ultimately save substantial time, funds and effort while producing more reliable and repeatable population counts, which are crucially important to monitor wildlife populations (Torney et al., 2019). Here we applied the approach to a video that was obtained opportunistically and generated for a documentary film; thus, filming decisions such as altitude and angle were made to maximize visual impact, rather than to facilitate accurate counting of individual animals. The small size of the animals in the video and the high degree of occlusion of animals by conspecifics (due to the oblique camera angle) complicated the detection and tracking tasks and required more manual correction effort than would be likely necessary if our approach were applied to videos captured for the purposes of counting animals. Because any future videos collected explicitly for counting purposes are expected to differ substantially from the video used in this study, we decided to forgo significant refinement of our detection and tracking procedures. For future mega-herd counts, we would aim to establish a standardized filming protocol that would include using an overhead (nadir) filming angle to reduce occlusions, flying within a consistent altitude range, and filming in the middle of the day (if compatible with targeted buffalo behaviour) or under cloudy conditions to reduce shadows (young buffalo can be difficult to identify in the shadow of their mothers). This would generate consistent high-quality imagery across filming campaigns, which would reduce the amount of retraining of the detection model required for subsequent counts. Such guidelines could be shared with wildlife filming companies to encourage them to contribute standardized footage to be used for monitoring and conservation purposes.

Aside from the unusually large number of individuals aggregating in the MD, the information provided by stakeholders aligned well with scientific understanding of Cape buffalo ecology. The Mababe mega-herd is most likely formed by several groups congregating on shared resources, representative of the documented Cape buffalo fission-fusion society whereby groups of individuals come together and separate depending on a variety of factors, such as forage, water and predation pressure (Cross et al., 2005; Taylor et al., 2023; Wielgus et al., 2020). The size of the mega-herd, combined with stakeholder assertions that the number of buffalo has not diminished over recent years and supported by aerial survey estimates, provides evidence that the MD ecosystem is maintaining its functionality, although accurate long-term monitoring of several species would be needed to confirm this (Kiffner et al., 2020). The flow of water into the MD during the hottest and driest time of the year, combined with productive grasses from lacustrine soils (Fynn et al., 2014), appears to be providing optimal conditions for the formation of this mega-herd. Stakeholders agreed that the mega-herd splits into smaller components during the rainy season, but opinions diverged on their rainy season ranges, indicating a need for further research on their movement patterns. Sianga et al., 2017 found that some buffalo occupying the Savuti Marsh, north of the MD, during the rainy season, moved east to the Linyanti Swamps during the dry season, but overall little is known about buffalo movements around the MD.

Reliable, accurate and repeatable counts are essential for population monitoring (Marsh & Trenham, 2008), and these can be related to the impacts of anthropogenic activities or shifts in environmental conditions. Cape buffalo are classified as Least Concern on the IUCN red list (Cornelis et al., 2023), but their high level of water-dependence makes them vulnerable to population declines in the hotter, drier conditions predicted by climate change in southern Africa (Engelbrecht et al., 2015). The strong seasonality of the presence of the Mababe megaherd in the MD highlights the importance of access to different habitats and the ability to move across landscapes for the conservation of large herbivores (OwenSmith et al., 2020). Repeated application of the novel counting method described here, as well as continued interaction with local stakeholders, could allow the detection of ecological disturbances (Service et al., 2014) that could affect buffalo and predict potential trophic cascades resulting from changes to the buffalo population. Successful conservation is boosted by a sense of environmental stewardship in local stakeholders (Adade Williams et al., 2020), who can promote sustainable practices and prevent illegal offtake. By encouraging stakeholders to report sightings through citizen science, their involvement in monitoring and conservation can be increased and could lead to the documentation of other unknown aggregations or unusual phenomena.

The Mababe mega-herd is likely to have knock-on ecological impacts for other species, as suggested by some stakeholders. A combination of trampling and heavy grazing pressure could form a basis for facilitation of smaller grazers; further research into vegetation characteristics and herbivore interactions would be beneficial in elucidating such processes. The mega-herd may be supporting unusually high populations of non-pride-holding male lions, which could be very successful at hunting weaker individuals on the periphery of the megaherd. As large-bodied herbivores, buffalo face a higher extinction risk than smaller-bodied species (Di Marco et al., 2014), and a decline in buffalo numbers could affect predators and sympatric grazing species. Buffalo, as one of the Big Five African species, are sought after on safari, particularly when they form large groups, so the Mababe mega-herd also constitutes an ecological asset for stakeholders (Chardonnet et al., 2023). The lack of awareness of the mega-herd displayed by Mababe community members highlights the disparity in socioeconomic costs and benefits derived from natural resources by different stakeholder groups (Snyman, 2014). Mababe community members primarily experience buffalo as risks to be avoided, although some individuals are employed by local tourism operators. Other stakeholders were better informed about the mega-herd and had generally benefited more from the herd through profits derived from tourism or wildlife filmmaking. On the other hand, this lack of awareness in the community may also be interpreted as buffalo avoiding humans, their livestock, and settlements, as has been observed elsewhere (VallsFox et al., 2018). Given the potential role of buffalo in human-wildlife conflicts (e.g., crop destruction, harm to people, pathogen transmission), the coexistence of the largest recorded buffalo herd with human rural populations without major conflict is a positive sign.

## CONCLUSION

Despite an ever-growing human presence in and influence on natural systems, there are still some wildlife phenomena, such as the Mababe mega-herd of Cape buffalo, that have not yet been documented, even if these are known to local stakeholders. Opportunistic material collection can be a useful source of ecological knowledge even if not produced using a strict scientific approach. Evidence from historic and current scientific surveys and LEK indicate that the Mababe mega-herd has been maintaining or possibly increasing its size over the last decade, suggesting a healthy ecosystem. Communication among people with different areas of expertise can lead to collaborative opportunities to apply technology to conservation challenges, supporting the acquisition of information vital for population monitoring. The initial development of the software highlighted here requires ground-truthing and training, so can be a relatively lengthy process, but the long-term benefits will outweigh short-term costs, given the accuracy of the method and its potential for replicability under similar conditions. Accurate counts should be combined with LEK from stakeholders to add ecological context that can inform conservation efforts, and non-scientific sources of information should be acknowledged as being able to produce valuable insights. Expanding the scope of the approaches presented here to other species and situations will further increase the benefits of such applications.

## DATA ACCESSIBILITY

The data used in this submission have been uploaded to the Edmond data repository and are accessible for peer review through this link: https://edmond.mpg.de/privateurl.xhtml?token=233f85da-c41f-481f-9bc3-bceb313227a8. Upon acceptance of this manuscript, the metadata for the data repository will be updated to reference the published version of the manuscript, and the data repository will be published and be permanently accessible through a DOI.

The original video clip used for the analysis is the commercial property of Netflix and is therefore not included in the linked data repository and cannot be made publicly available. However, all elements derived from the original clip (i.e. cropped video frames) for analysis are included in the data repository, enabling the replication of the results reported in this manuscript. This submission uses a combination of novel and published code. The novel code is available in a GitHub repository accessible through this link: https://github.com/BCostelloe/mababe-megaherd. The ReadMe document of the repository includes links to published code and clear references to the corresponding papers where appropriate. In the manuscript, descriptions of analyses using published code also include clear references to the original publication. Upon acceptance of this manuscript, we will publish a release of this repository so that the accepted version of the code remains accessible indefinitely.

## ACKNOWLEDGMENTS

The authors thank Silverback Productions and Netflix for supplying the documentary footage. We thank all informants for their valuable information: Brad Bestelink, Cobus Calitz, Mike Holding and Tshiamo Kebuelemang. B.R.C. acknowledges support from the Deutsche Forschungsgemeinschaft (DFG, German Research Foundation) under Germany’s Excellence Strategy - Centre for the Advanced Study of Collective Behaviour EXC 2117-422037984 and support from NVIDIA Corporation’s Academic Hardware Grant Program.

## AUTHOR CONTRIBUTIONS

Study conceptualization: AC, EB, BRC; Data collection: EB, GM; Data analysis: BRC, BK; Manuscript preparation: EB, BRC; Manuscript editing: EB, BRC, BK, EW, GM, AC.

## COMPETING INTERESTS

The authors declare no competing interests

## Notes

https://edmond.mpg.de/privateurl.xhtml?token=233f85da-c41f-481f-9bc3-bceb313227a8

https://github.com/BCostelloe/mababe-megaherd

### Competing Interest Statement

The authors have declared no competing interest.

