## Supplemental Text for "Combining artificial intelligence and local ecological knowledge to document the largest ever-counted Cape buffalo mega-herd"

### I. DETECTION MODEL PERFORMANCE

We evaluated the performance of our buffalo detection model using the images in the validation set as a ground truth. We calculated average precision (AP) using an Intersection over Union (IoU) value of 0.5. AP measures the models ability to correctly detect buffalo, accounting for both the precision of the model (i.e. the proportion of predictions that correspond to actual buffalo) and its recall (i.e. the proportion of actual buffalo that the model detected). AP values can range between 0 and 1, with 1 indicating perfect model performance. The IoU value provides a measure of accuracy of individual detections by comparing the bounding boxes drawn by human annotators to those generated by the detection model: the intersection (area of overlap) of the two boxes is divided by the union (the total combined area of the two boxes) to generate an IoU value between 0 (no overlap) and 1 (perfect overlap).

When applied to the validation images, the detection model had an AP of 0.868, indicating that on average 86.8% of buffalo were detected correctly with bounding boxes that overlapped the human-generated bounding boxes by at least 50%. Figure S1 shows model performance at a range of IoU values. Model performance is very good at low IoU values but declines drastically at higher IoU values. For our purposes, it is not important that the bounding boxes generated by the model align precisely with the extent of the buffalo. For downstream tasks (tracking and manual review), bounding boxes are converted into points by taking their centroids; therefore the boxes themselves must only be accurate enough that their centroids fall somewhere on the detected buffalos body. We therefore accepted this model performance as sufficient.

### II. MANUAL REVIEW OF TRACKED VIDEOS

Table S1 gives the results for the first round of manual review, in which each automatically tracked video clip was reviewed independently by 3 reviewers (BRC, AC, EW). If all three reviewers agreed on the total count, the count was accepted. The sum of counts from these clips is $R_1$ in Equation 3 in the Main Text. If the reviewers disagreed on the count for a given clip, the clip was re-reviewed in the second round of review.

Table S2 gives the results of the second round of review, in which one reviewer (BRC) reviewed the 34 clips not accepted in Round 1, and a second reviewer (BK) reviewed a randomly-selected subset of 8 clips for the purposes of quantifying BRCs detection probability and precision. Each reviewer was randomly assigned as the primary observer for four clips.

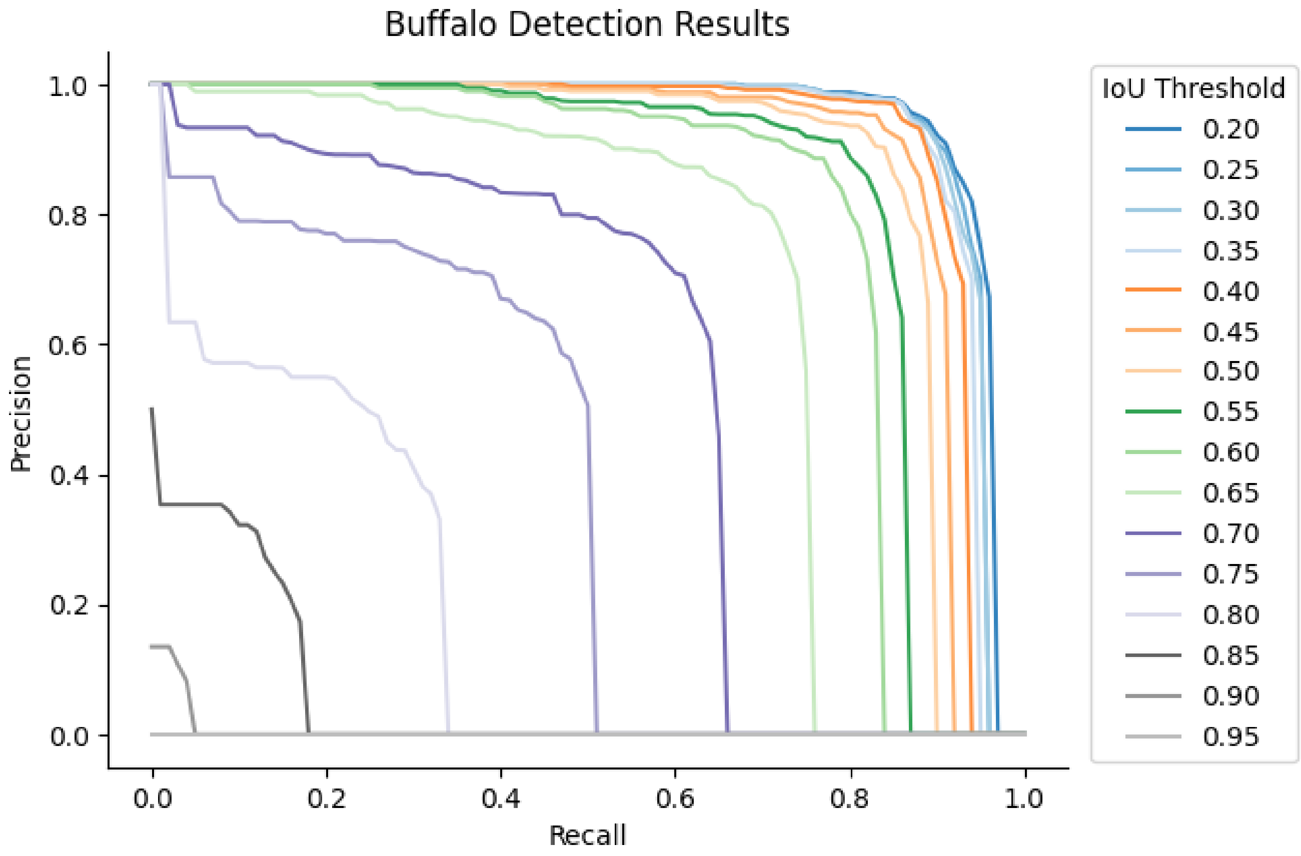


**Figure S1:** Precision-recall curves illustrating the performance of the detection model. Precision indicates the proportion of the models predictions that correspond to actual buffalo. Recall indicates the proportion of buffalo that were detected by the model. Each curve corresponds to a different IoU threshold. Higher IoU thresholds only consider detections with higher overlap between manually-annotated bounding boxes and those predicted by the model; lower IoU thresholds relax this criterion. The AP value reported in the main text was generated using an IoU threshold of 0.5.

**Table S1:** Results of the first round of manual count review. Shaded rows are video clips for which all reviewers agreed. Note that the count for output_video_0_18 includes 15 buffalo that were visible in the video, but had not yet crossed the count line before the video ended.

| | Counts | | | |
|---|---|---|---|---|
| **Clip name** | **BRC** | **AC** | **EW** | **Accepted (Round 1)** |
| output_video_0_0 | 0 | 0 | 0 | 0 |
| output_video_0_1 | 0 | 0 | 0 | 0 |
| output_video_0_2 | 0 | 0 | 0 | 0 |
| output_video_0_3 | 0 | 0 | 0 | 0 |
| output_video_0_4 | 0 | 0 | 0 | 0 |
| output_video_0_5 | 0 | 0 | 0 | 0 |
| output_video_0_6 | 0 | 0 | 0 | 0 |
| output_video_0_7 | 0 | 0 | 0 | 0 |
| output_video_0_8 | 0 | 0 | 0 | 0 |
| output_video_0_9 | 0 | 0 | 0 | 0 |

| | Counts | | | |
|---|---|---|---|---|
| Clip name | BRC | AC | EW | Accepted (Round 1) |
| output_video_0_10 | 1 | 1 | 1 | 1 |
| output_video_0_11 | 0 | 0 | 0 | 0 |
| output_video_0_12 | 9 | 8 | 7 | NA |
| output_video_0_13 | 1 | 1 | 1 | 1 |
| output_video_0_14 | 3 | 3 | 3 | 3 |
| output_video_0_15 | 4 | 6 | 3 | NA |
| output_video_0_16 | 3 | 3 | 2 | NA |
| output_video_0_17 | 8 | 11 | 9 | NA |
| output_video_0_18 | 19 | 19 | 19 | NA |
| output_video_1_0 | 0 | 0 | 0 | 0 |
| output_video_1_1 | 0 | 0 | 0 | 0 |
| output_video_1_2 | 1 | 1 | 1 | 1 |
| output_video_1_3 | 9 | 9 | 9 | 9 |
| output_video_1_4 | 17 | 18 | 16 | NA |
| output_video_1_5 | 17 | 18 | 18 | NA |
| output_video_1_6 | 15 | 15 | 14 | NA |
| output_video_1_7 | 9 | 10 | 9 | NA |
| output_video_1_8 | 9 | 8 | 9 | NA |
| output_video_1_9 | 15 | 16 | 16 | NA |
| output_video_1_10 | 24 | 21 | 23 | NA |
| output_video_1_11 | 17 | 13 | 18 | NA |
| output_video_1_12 | 28 | 24 | 28 | NA |
| output_video_1_13 | 6 | 6 | 6 | 6 |
| output_video_1_14 | 18 | 17 | 19 | NA |
| output_video_1_15 | 9 | 8 | 9 | NA |
| output_video_1_16 | 13 | 14 | 11 | NA |
| output_video_1_17 | 3 | 9 | 3 | NA |
| output_video_1_18 | 0 | 0 | 0 | 0 |
| output_video_2_0 | 0 | 0 | 0 | 0 |
| output_video_2_1 | 11 | 12 | 11 | NA |
| output_video_2_2 | 90 | 82 | 82 | NA |
| output_video_2_3 | 42 | 40 | 42 | NA |
| output_video_2_4 | 10 | 8 | 9 | NA |
| output_video_2_5 | 6 | 7 | 9 | NA |

| | Counts | | | |
|---|---|---|---|---|
| Clip name | BRC | AC | EW | Accepted (Round 1) |
| output_video_2_6 | 3 | 6 | 3 | NA |
| output_video_2_7 | 2 | 1 | 2 | NA |
| output_video_2_8 | 3 | 3 | 3 | 3 |
| output_video_2_9 | 3 | 2 | 3 | NA |
| output_video_2_10 | 1 | 1 | 1 | 1 |
| output_video_2_11 | 4 | 5 | 4 | NA |
| output_video_2_12 | 5 | 5 | 5 | 5 |
| output_video_2_13 | 0 | 0 | 0 | 0 |
| output_video_2_14 | 0 | 1 | 0 | NA |
| output_video_2_15 | 0 | 0 | 0 | 0 |
| output_video_2_16 | 0 | 0 | 0 | 0 |
| output_video_2_17 | 0 | 0 | 0 | 0 |
| output_video_2_18 | 0 | 0 | 0 | 0 |
| output_video_3_0 | 0 | 0 | 0 | 0 |
| output_video_3_1 | 95 | 91 | 95 | NA |
| output_video_3_2 | 101 | 100 | 95 | NA |
| output_video_3_3 | 17 | 16 | 19 | NA |
| output_video_3_4 | 0 | 0 | 0 | 0 |
| output_video_3_5 | 0 | 0 | 0 | 0 |
| output_video_3_6 | 0 | 0 | 0 | 0 |
| output_video_3_7 | 0 | 0 | 0 | 0 |
| output_video_3_8 | 0 | 0 | 0 | 0 |
| output_video_3_9 | 0 | 0 | 0 | 0 |
| output_video_3_10 | 0 | 0 | 0 | 0 |
| output_video_3_11 | 0 | 0 | 0 | 0 |
| output_video_3_12 | 0 | 0 | 0 | 0 |
| output_video_3_13 | 0 | 0 | 0 | 0 |
| output_video_3_14 | 0 | 0 | 0 | 0 |
| output_video_3_15 | 0 | 0 | 0 | 0 |
| output_video_3_16 | 0 | 0 | 0 | 0 |
| output_video_3_17 | 0 | 0 | 0 | 0 |
| output_video_3_18 | 0 | 0 | 0 | 0 |
| output_video_4_0 | 0 | 0 | 0 | 0 |
| output_video_4_1 | 12 | 11 | 12 | NA |

| | Counts | | | |
|---|---|---|---|---|
| Clip name | BRC | AC | EW | Accepted (Round 1) |
| output_video_4_2 | 22 | 25 | 21 | NA |
| output_video_4_3 | 3 | 3 | 2 | NA |
| output_video_4_4 | 0 | 0 | 0 | 0 |
| output_video_4_5 | 0 | 0 | 0 | 0 |
| output_video_4_6 | 0 | 0 | 0 | 0 |
| output_video_4_7 | 0 | 0 | 0 | 0 |
| output_video_4_8 | 0 | 0 | 0 | 0 |
| output_video_4_9 | 0 | 0 | 0 | 0 |
| output_video_4_10 | 0 | 0 | 0 | 0 |
| output_video_4_11 | 0 | 0 | 0 | 0 |
| output_video_4_12 | 0 | 0 | 0 | 0 |
| output_video_4_13 | 0 | 0 | 0 | 0 |
| output_video_4_14 | 0 | 0 | 0 | 0 |
| output_video_4_15 | 0 | 0 | 0 | 0 |
| output_video_4_16 | 0 | 0 | 0 | 0 |
| output_video_4_17 | 0 | 0 | 0 | 0 |
| output_video_4_18 | 0 | 0 | 0 | 0 |
| **Total** | **688** | **678** | **672** | **30** |

**Table S2:** Results of the second round of manual count review.

| Clip name | BRC | BK | Primary observer |
|---|---|---|---|
| output_video_0_12 | 9 | | |
| output_video_0_15 | 3 | 3 | BRC |
| output_video_0_16 | 3 | | |
| output_video_0_17 | 11 | | |
| output_video_0_18 | 19 | | |
| output_video_1_4 | 18 | 18 | BRC |
| output_video_1_5 | 18 | | |
| output_video_1_6 | 14 | | |
| output_video_1_7 | 9 | | |
| output_video_1_8 | 9 | | |
| output_video_1_9 | 17 | 16 | BK |
| output_video_1_10 | 25 | | |
| output_video_1_11 | 17 | 17 | BRC |

| Clip name | BRC | BK | Primary observer |
|---|---|---|---|
| output_video_1_12 | 29 | 28 | BK |
| output_video_1_14 | 19 | | |
| output_video_1_15 | 9 | | |
| output_video_1_16 | 11 | | |
| output_video_1_17 | 3 | 3 | BRC |
| output_video_2_1 | 11 | | |
| output_video_2_2 | 9 | | |
| output_video_2_3 | 45 | | |
| output_video_2_4 | 9 | | |
| output_video_2_5 | 9 | | |
| output_video_2_6 | 3 | | |
| output_video_2_7 | 2 | | |
| output_video_2_9 | 3 | | |
| output_video_2_11 | 4 | 3 | BK |
| output_video_2_14 | 0 | | |
| output_video_3_1 | 103 | | |
| output_video_3_2 | 102 | | |
| output_video_3_3 | 20 | | |
| output_video_4_1 | 12 | 11 | BK |
| output_video_4_2 | 21 | | |
| output_video_4_3 | 3 | | |
| **Total** | **681** | | |

### III. ESTIMATING THE NUMBER OF BUFFALO MISSED DUE TO VIDEO ENDING PREMATURELY

To estimate the rates of buffalo passing the count line beyond the duration of the video, created time series of our count data and plotted these using bins of different time durations (2, 5 and 10 seconds). We fit a Bayesian Poisson regression model with a Gaussian-shaped time-varying intensity function to each time series, and used these to predict rates of buffalo crossing for future time bins. We defined cut-off threshold rates of 1 buffalo crossing the line every 5, 10 and 20 seconds, or 0.2, 0.1 and 0.05 buffalo per second. These rates are relative to real time, not the timeframe of the sped-up video. Figure S2 shows the plotted data, fit lines, and cut-off thresholds for each time series. Table S3 gives the predict herd duration beyond the end of the video, and Table S4 gives the estimated cumulative number of buffalo that were missed by the video under each simulated scenario.

| | | Cut-off threshold (buffalo crossings per second) | | |
|---|---|---|---|---|
| | | **0.2** | **0.1** | **0.05** |
| **Bin size (seconds)** | **2** | 332 (1207.5) | 446 (1672.5) | 446 (1672.5) |
| | **5** | 285(1068.75) | 380 (1425) | 500 (1875) |
| | **10** | 290 (1087.5) | 360 (1350) | 450 (1687.5) |

**Table S3:** Predicted herd crossing durations beyond the end of the video, under predictions generated using different time bin sizes and cut-off thresholds. Values reported are in seconds and on the time scale of the sped-up video; real time durations are given in parentheses.

| | | Cut-off threshold (buffalo crossings per second) | | |
|---|---|---|---|---|
| | | **0.2** | **0.1** | **0.05** |
| **Bin size (seconds)** | **2** | 471 [69, 941] | 481 [67, 1003] | 481 [67, 1003] |
| | **5** | 442 [152, 724] | 455 [152, 770] | 459 [152, 794] |
| | **10** | 468 [231, 697] | 479 [231, 731] | 485 [231, 752] |

**Table S4:** . Cumulative buffalo missed due to early termination of the video, under predictions generated using different time bin sizes and cut-off thresholds. Values reported are the mean prediction with the 95% confidence range given in brackets.

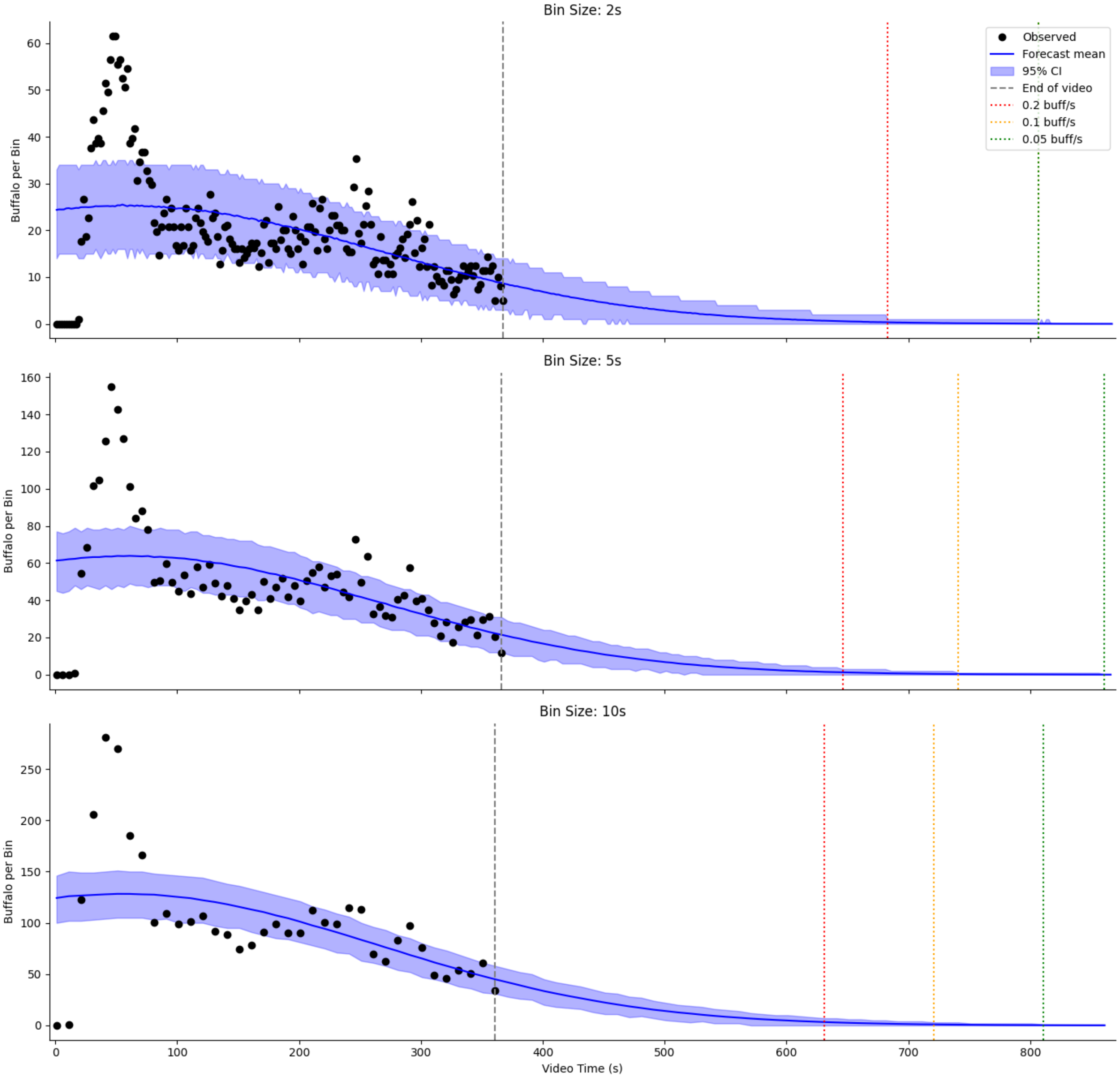


**Figure S2:** Buffalo crossing forecasts. Data points represent the number of buffalo that cross the count line in each time bin, including automatically tracked and manually counted buffalo. The herd is assumed to end when the upper 95% confidence interval crosses a threshold rate (0.2, 0.1 or 0.05 buffalo crossing per second, indicated by colored vertical lines). The vertical grey line indicates the end of the video.
